# Post-weaning social isolation increases ΔFosB/FosB protein expression in the prefrontal cortex and hippocampus in mice

**DOI:** 10.1101/2020.06.04.135517

**Authors:** Michael Noback, Gongliang Zhang, Noelle White, James C. Barrow, Gregory V. Carr

## Abstract

Social isolation is a growing public health concern across the lifespan. Specifically, isolation early in life, during critical periods of brain development, increases the risk of psychiatric disorders later in life. Previous studies of isolation models in mice have shown distinct neurological abnormalities in various regions of the brain, but the mechanism linking the experience of isolation to these phenotypes is unclear. In this study, we show that ΔFosB, a long-lived transcription factor associated with chronic stress responses and drug-induced neuroplasticity, is upregulated in the medial prefrontal cortex and hippocampus of adult C57BL/6J mice isolated for two weeks post-weaning. Additionally, a related transcription factor, FosB, is also increased in the medial prefrontal cortex in socially isolated females. These results show that short-term isolation during the critical post-weaning period has long-lasting and sex-dependent effects on gene expression in brain.

## 1. Introduction

Social isolation (SI) during childhood and adolescence is an adverse event that increases the risk for developing several psychiatric disorders later in life, including anxiety, depression, and schizophrenia [1]. SI has long been recognized as a public health issue among older populations [2], but recent evidence reveals it is a growing problem among adolescents and young adults [3]. The potential disruption of critical developmental processes compounds the negative consequences of SI for this age group. Despite the wealth of epidemiological data available on the detrimental effects of SI, little is known about the underlying neural mechanisms through which SI increases risk for psychiatric disorders [1].

Some of the immediate effects of SI include impaired cognition, increased risk for substance use disorders, and increased anxiety and depression. Some of these effects are also observed in situations where stress response proteins are dysregulated. The transcription factor FosB is transiently expressed in response to environmental stress, but in situations of persistent stress, the truncated and longer-lived isoform ΔFosB tends to accumulate [4]. High levels of ΔFosB are observed in response to chronic drug exposure and are associated with addiction-like behaviors in rodents [5]. Increases in ΔFosB are also observed in chronic social stress models, specifically in repeated social defeat stress [6].

In social defeat animal models, ΔFosB elevation is observed in the medial prefrontal cortex (mPFC), a phenomenon associated with increased susceptibility to social stress, and increased anxiety- and depression-like behaviors. Viral-mediated genetic overexpression of ΔFosB produces a similar behavioral profile, which suggests that the transcription factor plays a causal role in the phenotype.

ΔFosB/FosB induction has been studied in the context of rodent SI models, but, to our knowledge, not in models of post-weaning/adolescent isolation [7,8]. Here we assessed the expression of ΔFosB/FosB in the mPFC, hippocampus, and striatum, three regions involved in the behavioral response to stress, after exposure to a validated model of post-weaning isolation [9–11].

## 2. Materials and methods

### 2.1. Mice

Male and female C57BL/6J mice (6-7 weeks old) were purchased from The Jackson Laboratory (Bar Harbor, ME, USA) and set up as breeding pairs. Only one litter from each breeding pair was used in these experiments. The isolation procedure used for these studies was adapted from the procedure described by Makinodan and colleagues [9]. Briefly, pups were weaned at postnatal day 21 (P21) and either housed in same sex groups of three for the duration of the experiment or singly housed from P21-P35 and then rehoused with another isolated littermate for the duration of the experiment (Figure 1).

**Figure 1.**
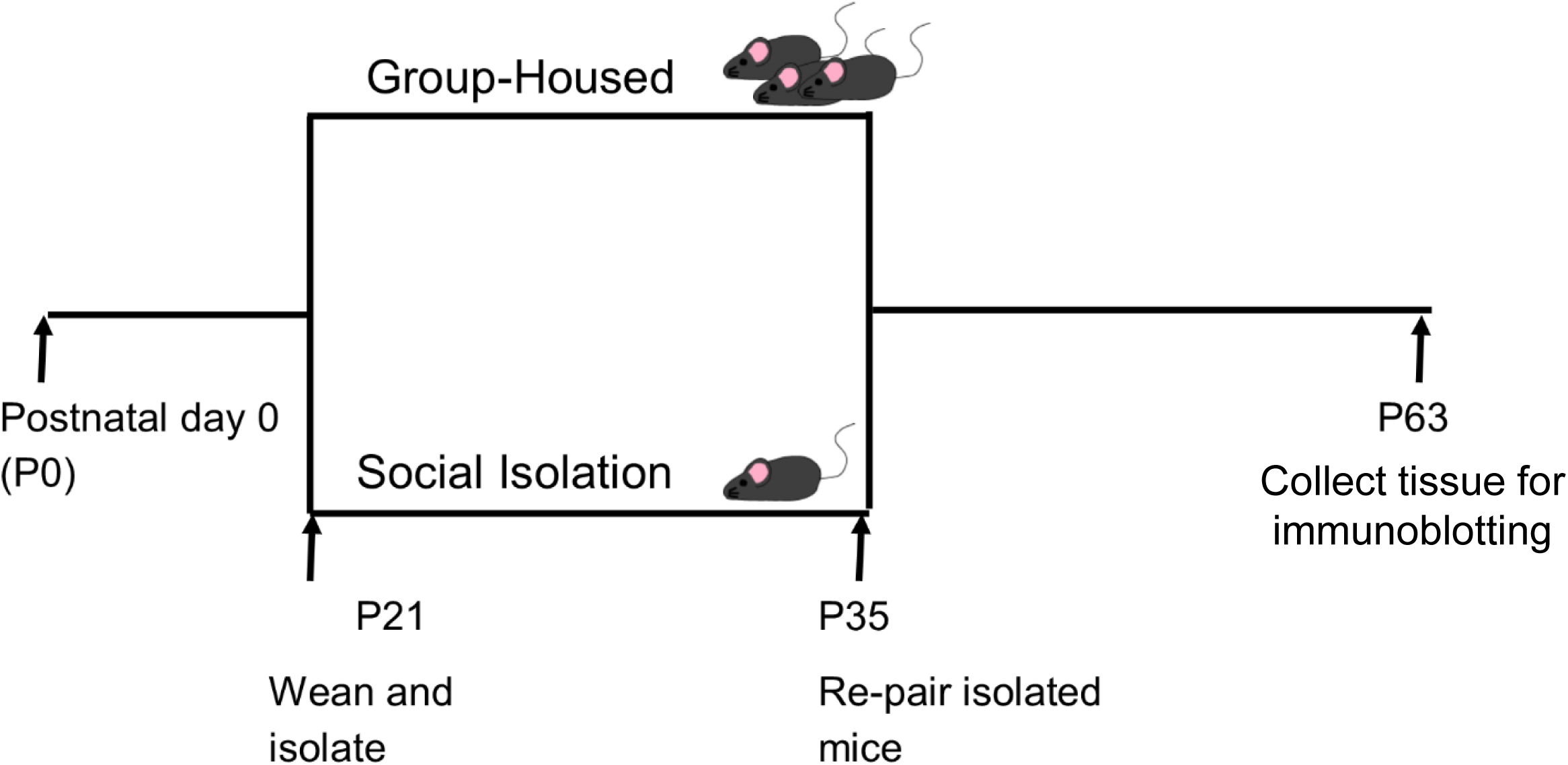
Post-weaning isolation protocol. Mice are weaned at P21 into group-housed cages (3 mice/cage) or isolation cages. On P35, isolated mice are repaired with previously isolated littermates. Tissue was collected at P63.

### 2.2. Immunoblotting

At P63, mice were anesthetized with isoflurane and then decapitated. The brain was removed and the mPFC (infralimbic and prelimbic subregions), hippocampus, and striatum were dissected using previously described methods [12] and flash frozen on dry ice and stored at -80°C until processing. The tissue was homogenized and sonicated in T-Per lysis buffer (Thermo Scientific, Rockford, IL, United States). The protein concentration of the samples was determined using a Pierce BCA Protein Assay Kit (Thermo Scientific, 23225). 40 µg of protein was separated by NuPAGE 4-12% Bis-Tris Protein Gels (Invitrogen, NP0335BOX) and transferred to a nitrocellulose membrane with an iBlot® Transfer Stack (ThermoFisher Scientific, IB301001). After blockade with Odyssey blocking buffer (LI-COR, Lincoln, NE, USA; 927-40000) for 1 h at room temperature, the membrane was incubated with the rabbit monoclonal anti-FosB primary antibody (dilution: 1:1000, product number: 2251S, Cell Signaling Technology, Danver, MA, USA) at 4 °C overnight. ΔFosB/FosB levels were normalized to beta-actin, so membranes were also incubated with a mouse monoclonal anti-beta actin primary antibody (dilution: 1:5000, product number: ab8226, Abcam, Cambridge, MA, USA). After TBST washing three times for 10 minutes each wash, the membrane was incubated with the corresponding secondary antibodies (goat anti-rabbit IRDye^®^ 800CW and goat anti-mouse IRDye^®^ 680LT (dilutions: 1:15,000, LI-COR) for 1 h at room temperature. The western blot protein bands were captured by Odyssey CLX and analyzed by Image Studio software (V3.1, LI-COR).

## 3. Results

### 3.1. ΔFosB protein is elevated in the mPFC and hippocampus following SI stress

We found that ΔFosB protein levels were increased in the mPFC (*F*_*1,33*_ = 16.54, *p* = 0.0003) and hippocampus (*F*_*1,33*_ = 4.666, *p* = 0.0381) of adult mice exposed to post-weaning SI compared to group-housed littermates. There were no effects of sex on ΔFosB levels in either region (mPFC: *F*_*1,33*_ = 0.0270, *p* = 0.8704; Hippocampus: *F*_*1,33*_ = 0.7638, *p* = 0.3885) or sex X housing interactions (mPFC: *F*_*1,33*_ = 0.0270, *p* = 0.8704; Hippocampus: *F*_*1,33*_ = 0.9484, *p* = 0.3372). There were no significant effects of sex (*F*_*1,33*_ = 0.0986, *p* = 0.7555), housing (*F*_*1,33*_ = 1.427, *p* = 0.2408), or sex X housing interactions (*F*_*1,33*_ = 0.0986, *p* = 0.7555) on ΔFosB in the striatum (Figure 2).

**Figure 2.**
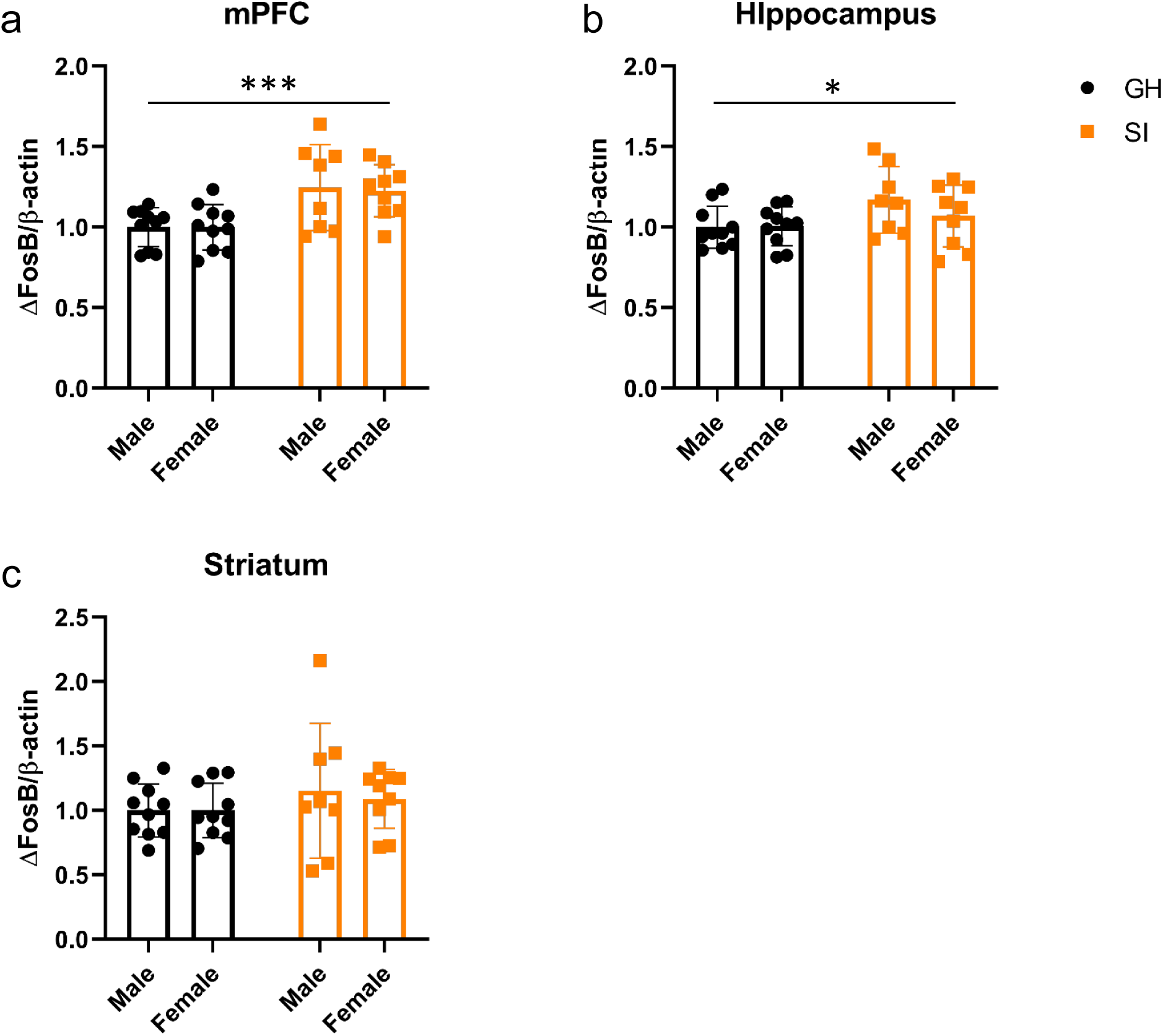
Relative ΔFosB protein levels. SI increases ΔFosB protein in the mPFC (a) and hippocampus (b) of male and female mice compared to their group-housed (GH) littermates. There were no significant differences in the striatum (c). n = 10/GH males, 10/GH females, 8/SI males, and 9/SI females. Data are normalized to the GH male group and represent the mean ± SEM. \**p < 0.05;* \*\*\**p <* 0.001.

### 3.2. FosB protein is decreased in male mice exposed to SI stress

We also measured the amount of FosB protein in the same three brain regions. There was a significant interaction between sex and housing on FosB in the mPFC (*F*_*1,33*_ = 5.708, *p* = 0.0228). *Post hoc* analyses showed that male mice exposed to SI have less FosB compared to female mice exposed to SI (*p* = 0.0160). There were no other significant pairwise differences between the groups. There were no significant effects of sex, housing, or sex X housing interactions in the hippocampus or striatum (Figure 3).

**Figure 3.**
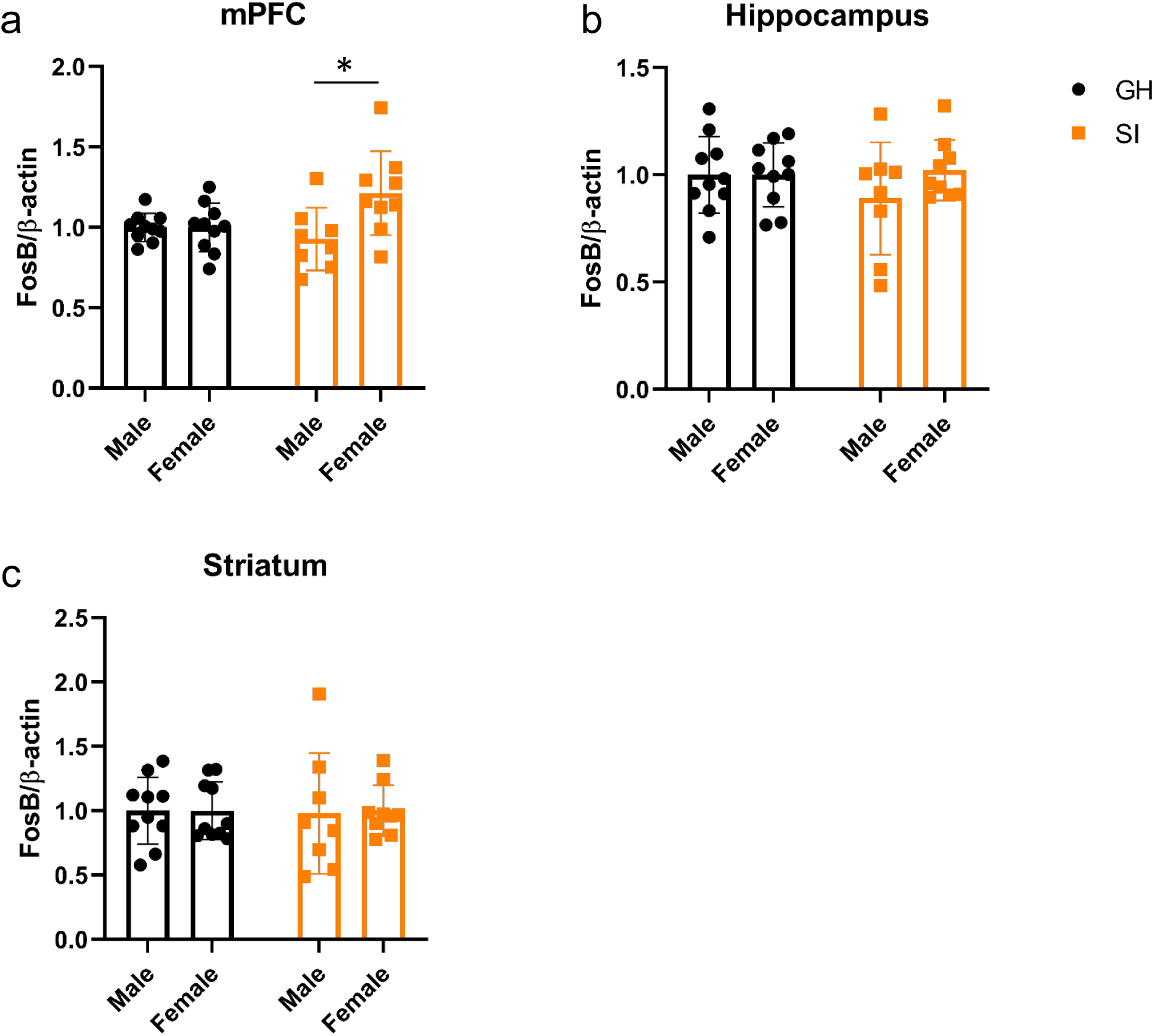
Relative FosB protein levels. SI increases FosB protein in the mPFC (a) of female mice compared to SI male mice. There were no other significant differences in the mPFC, hippocampus (b), or striatum (c). n = 10/GH males, 10/GH females, 8/SI males, and 9/SI females. Data are normalized to the GH male group and represent the mean ± SEM. \**p < 0.05*.

### 3.3. ΔFosB/FosB ratio is higher in the mPFC of male mice exposed to SI stress

Due to the changes in ΔFosB and FosB protein levels across multiple regions, we also analyzed the relative changes in these two proteins within individual mice. We found a significant interaction between sex and housing on ΔFosB/FosB ratio in the mPFC (*F*_*1,33*_= 8.585, *p* = 0.0061). *Post hoc* analyses showed that the interaction was driven by an increase in the ratio in SI males compared to all of the other groups (*p* = 0.0017 compared to group-housed males and *p* < 0.0001 compared to both group-housed females and SI females; Figure 4).

**Figure 4.**
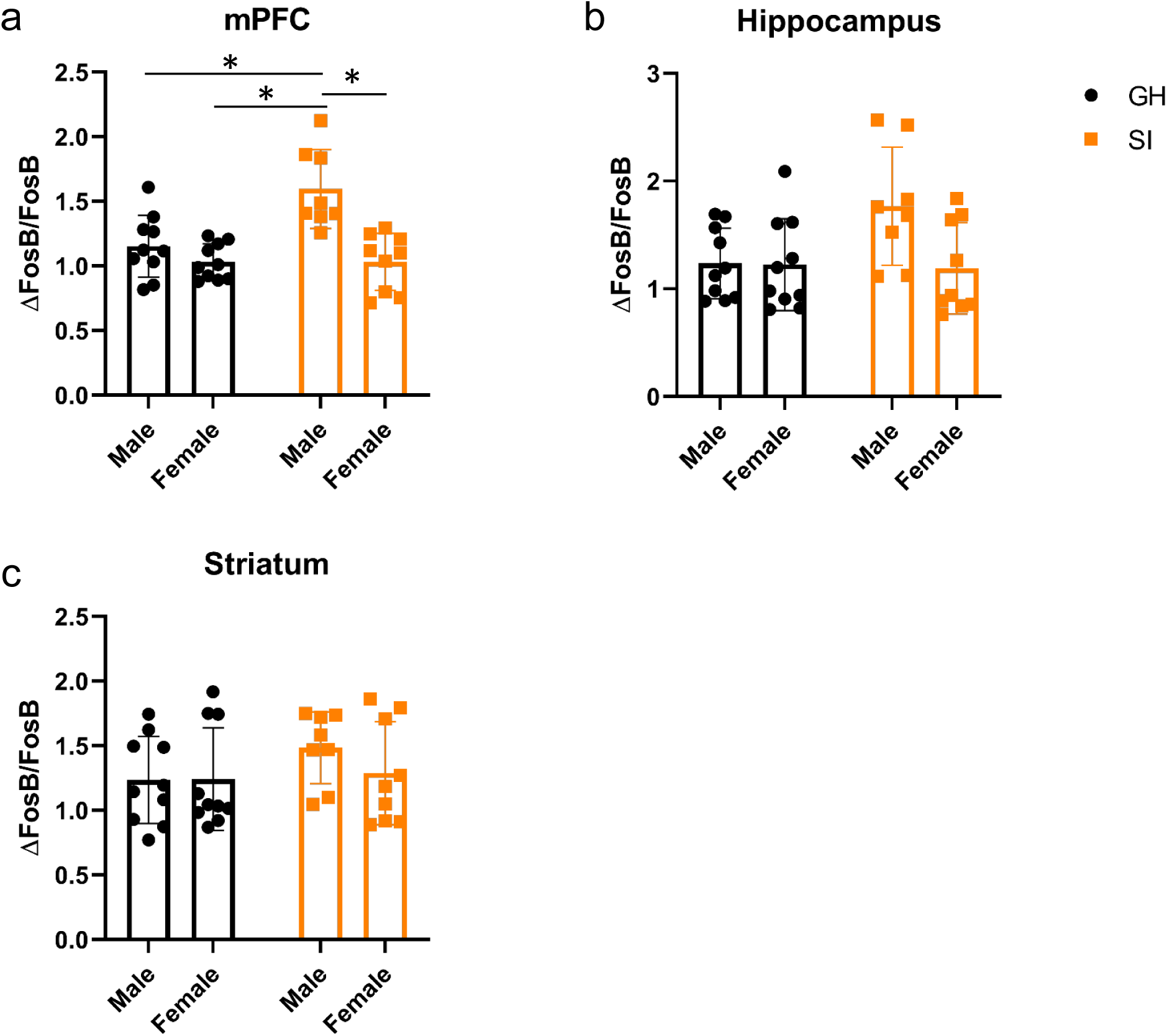
ΔFosB / FosB ratio. SI increases the ΔFosB / FosB ratio in the mPFC of SI male mice compared to the other three groups. There were no other significant differences in the mPFC, hippocampus (b), or striatum (c). n = 10/GH males, 10/GH females, 8/SI males, and 9/SI females. Data are the mean ± SEM. \**p < 0.05*.

## 4. Discussion

In this experiment, we measured the levels of ΔFosB and FosB protein in the mPFC, hippocampus, and striatum of mice exposed to transient post-weaning SI. We found that ΔFosB protein expression is a long-term marker of SI in this model. Specifically, ΔFosB protein is increased in the mPFC and hippocampus of adult mice (P63) that were socially isolated from P21-P35. Additionally, there were sex X housing interactions. In the mPFC, SI females had more FosB protein than SI males, but not group-housed females or males. We also measured the ΔFosB/FosB ratio in the three brain regions and found that SI male mice had more ΔFosB relative to FosB in the mPFC compared to the three other groups. Interestingly, these changes in ΔFosB/FosB expression are present weeks after the termination of the SI stress, suggesting there are long-lasting effects of SI that are not mitigated by a return to group-housing.

Increased ΔFosB protein is a hallmark of exposure to multiple types of stress. For example, ΔFosB levels are elevated following exposure to chronic, but not acute administration of multiple drugs of abuse in rats and mice [13]. Chronic restraint and unpredictable stress increase ΔFosB expression in the brain as well [14]. Moreover, chronic exposure to seemingly beneficial perturbations including wheel-running and antidepressants increase ΔFosB expression as well [15,16].

ΔFosB expression, particularly in the nucleus accumbens, is associated with augmented responses to drugs of abuse. Early experiments utilizing overexpression of ΔFosB indicated that high levels of the transcription factor in the nucleus accumbens increases the responsiveness of mice to the rewarding and locomotor-activating effects of cocaine [17]. Additionally, ΔFosB is associated with increased self-administration of cocaine and inhibition of the aversive effects of kappa opioid receptor activation [18,19]. Taken together, the combined effects of increased reward sensitivity and decreased aversion could support increased drug-seeking and susceptibility to addiction. The increase in drug-seeking associated with ΔFosB expression appears to be a general effect because it is seen with multiple drugs of abuse, including opioids [20].

To our knowledge, our study is the first to investigate changes in ΔFosB/FosB following transient post-weaning isolation, but there have been reports in other models of social isolation. Isolation for eight weeks on adult female prairie voles increased ΔFosB/FosB immunohistochemistry in the basolateral amygdala [8]. Interestingly, prolonged social isolation in adult mice decreases ΔFosB in the nucleus accumbens and increases susceptibility to the detrimental effects of social defeat stress [7]. We found no change in ΔFosB or FosB protein in the striatum. The discrepancy may be due to our sampling of the entire striatum, including both dorsal and ventral (nucleus accumbens) subregions, and/or the difference in the age of the mice during isolation. There may be fundamental differences in the long-term effects of SI depending on when the isolation occurs.

We found significant changes in ΔFosB/FosB protein in the mPFC. The post-weaning period we examined is critical for the development of the prefrontal cortex in rodents [21,22]. Chronic administration of drugs of abuse have been shown to increase ΔFosB in the PFC [13]. Additionally, chronic treatment with the antipsychotic haloperidol increases ΔFosB in the PFC and the increase in ΔFosB is associated with cognitive disruption [23]. Chronic social defeat stress also increases ΔFosB in the mPFC [24]. These data suggest that the increased ΔFosB produced by social isolation stress may be associated with detrimental behavioral effects in these mice.

Female mice exposed to isolation also had elevated levels of FosB. Most studies investigating ΔFosB/FosB have utilized immunohistochemistry and antibodies that bind both ΔFosB and FosB, making it impossible to determine which variant was responsible for the signal. Here, we used western blotting techniques that allowed us to separate ΔFosB and FosB by size and quantify relative changes between the two variants [25]. It is unclear what the differential effects would be of having either elevated ΔFosB alone or in combination with FosB. If the induction of FosB is related to acute stress and ΔFosB is a marker of a previously terminated stressor, the different response between males and females may represent a critical difference in the downstream effects of social isolation. One study utilizing mutant mice with variable levels of ΔFosB and FosB indicates that relative expression patterns may produce differential behavioral effects. Specifically FosB can antagonize the effects of accumulated ΔFosB [26].

This study represents an initial characterization of the long-term effects of transient post-weaning SI. The role, if any, of ΔFosB/FosB in the behavioral or neurobiological alterations produced by this model are unknown [9,11,27]. Future studies will address whether expression of genes regulated by ΔFosB/FosB are also modulated by isolation and whether there is a causal relationship between ΔFosB/FosB activity and the SI behavioral phenotype.

## Abbreviations

SI: social isolation
mPFC: medial prefrontal cortex
P: postnatal day

## Data Availability Statement

The datasets generated by this project are available upon request.

## Author Contributions

**Michael Noback:** Conceptualization, Methodology, Writing-Original Draft, Writing-Review & Editing; **Gongliang Zhang:** Conceptualization, Methodology, Investigation, Formal Analysis, Writing-Review & Editing; **Noelle White:** Investigation, Writing-Review & Editing; **James C. Barrow:** Writing-Review & Editing, Supervision, Project Administration, Funding Acquisition; **Gregory V. Carr:** Conceptualization, Methodology, Formal Analysis, Validation, Writing-Original Draft, Writing-Review & Editing, Supervision, Project Administration.

## Funding

This work was funded by the Lieber Institute for Brain Development

## Acknowledgements

The authors thank Anna Kolobova for technical assistance.

## Notes

### Competing Interest Statement

The authors have declared no competing interest.

